# Predicting drug resistance in *M. tuberculosis* using a Long-term Recurrent Convolutional Network

**DOI:** 10.1101/2020.11.07.372136

**Authors:** Amir Hosein Safari, Nafiseh Sedaghat, Hooman Zabeti, Alpha Forna, Leonid Chindelevitch, Maxwell Libbrecht

## Abstract

**Motivation:** Drug resistance in Mycobacterium tuberculosis (MTB) is a growing threat to human health worldwide. One way to mitigate the risk of drug resistance is to enable clinicians to prescribe the right antibiotic drugs to each patient through methods that predict drug resistance in MTB using whole-genome sequencing (WGS) data. Existing machine learning methods for this task typically convert the WGS data from a given bacterial isolate into features corresponding to single-nucleotide polymorphisms (SNPs) or short sequence segments of a fixed length *K* (*K*-mers). Here, we introduce a gene burden-based method for predicting drug resistance in TB. We define one numerical feature per gene corresponding to the number of mutations in that gene in a given isolate. This representation greatly reduces the number of model parameters. We further propose a model architecture that considers both gene order and locality structure through a Long-term Recurrent Convolutional Network (LRCN) architecture, which combines convolutional and recurrent layers.

**Results:** We find that using these strategies yields a substantial, statistically significant improvement over state-of-the-art methods on a large dataset of *M. tuberculosis* isolates, and suggest that this improvement is driven by our method’s ability to account for the order of the genes in the genome and their organization into operons.

**Availability:** The implementations of our feature preprocessing pipeline^1^ and our LRCN model^2^ are publicly available, as is our complete dataset^3^.

**Supplementary information:** Additional data are available in the *Supplementary Materials* document^4^.

## 1 INTRODUCTION

Drug resistance is the phenomenon whereby an infectious organism (also known as a pathogen) develops resistance to the drugs that are commonly used in its treatment [41]. In this paper, our focus is on *Mycobacterium tuberculosis* (MTB), the etiological agent of tuberculosis (TB). It is the deadliest infectious disease today and is responsible for almost 2 million deaths every year among 10 million new cases [42]. The challenge of drug-resistant TB is not only a concern for low and middle-income countries, but also high-income countries [32]. The importance of drug-resistant TB—and other drug-resistant pathogens—is due to the fact that, without novel antimicrobial drugs, the total deaths due to drug resistance may exceed 10 million people a year by 2050, which is higher than the current annual mortality due to cancer [28].

One way to mitigate the risk of drug resistance is to carry out a drug susceptibility test (DST) by growing the bacterial isolate in the presence of different drugs and prescribing a regimen consisting of drugs the isolate is susceptible to. However, this approach is time-consuming and labour-intensive, so treatment is typically started before its results become available, potentially leading to poor outcomes.

The use of whole-genome sequencing (WGS) data makes it possible to identify drug resistance in hours rather than days. Prior methods for identifying drug resistance from WGS data can be divided into two categories. The first category consists of catalogue methods, which involve testing the WGS data of an isolate for the presence of known mutations associated with drug resistance. These mutations are primarily single nucleotide polymorphisms (SNPs), though they can also be insertions or deletions (indels) [2]. An isolate is then predicted to be resistant if it has one or more such mutations [1, 6, 13, 16, 37]. However, this approach often has poor predictive accuracy [33], especially for situations involving novel drug resistance mechanisms or resistance to untested or rarely-used drugs [17].

The second category consists of machine learning (ML) methods, which aim to to predict drug resistance by using models trained directly on paired WGS and DST data [3, 7, 10, 19, 20, 43, 45]. The current state-of-the-art models include the wide-n-deep neural network (WnD) [3], DeepAMR [43], KOVER [10], and gradient-boosted trees (GBTs) [7, 20]. While existing machine learning methods achieve better accuracy than catalogue methods, there remains much room for improvement, especially for the less commonly-used second and third-line drugs, and such an improvement in accuracy holds the promise of positively affecting clinical outcomes. We refer the reader to Supplementary Section 1 for an extensive review of existing ML methods for drug resistance prediction.

Existing ML methods typically convert their input WGS data into features that correspond to SNPs or short sequence segments of a fixed length *K* (*K*-mers). These representations do not necessarily take advantage of the organization of genome sequences into genes, despite the fact that genes are the unit of structure supporting the functional changes responsible for drug resistance.

Here, we introduce a gene-centric method for predicting drug resistance in TB. We define one feature per gene which represents the number of mutations in that gene in a given isolate. This representation greatly decreases the number of features—and therefore model parameters—relative to a SNP-based representation. To our knowledge, while gene-based statistical tests—known as gene burden tests—are sometimes used for microbial genome-wide association studies [8, 12], and have been used to represent rare variants in a previous ML method [3], this work is the first systematic use of such gene burden features for drug resistance prediction using ML methods. In contrast to previous practice, where only the mutations leading to a specific type of protein change such as deleterious or non-synonymous mutations contribute to the gene’s burden, our features count all the mutations found in a gene.

We further propose a model that accounts for both the order and the locality structure of genes through a Long-term Recurrent Convolutional Network (LRCN) architecture, which combines convolutional and recurrent layers. LRCNs have recently been successfully used in other fields such as computer vision [9, 25] but, to our knowledge, this is the first use of an LRCN in computational biology for a task that is not directly related to image processing. We also introduce a multi-task approach for this problem, where we train a single model to jointly predict resistance to twelve drugs [3, 44].

Remarkably, we find that using these strategies yields a substantial improvement over the state-of-the-art tools. This improvement is statistically significant and consistent across many drugs and settings, and requires both elements of our model, namely, the gene burden-based features and the LRCN model architecture. We verify via permutation testing that the order of genes and their organization into operons drive the model’s performance. Based on these results, we expect that this gene burden-based method may prove useful for other genotype-to-phenotype prediction problems.

## 2 MATERIALS AND METHODS

### 2.1 Data

To train and evaluate our method, we used the Pathosystems Resource Integration Center (PATRIC) [40] and the Relational Sequencing TB Data Platform (ReSeqTB) [36] datasets to collect 7,845 isolates, together with their resistance/susceptible status (labels) for twelve drugs. These include five first-line drugs—isoniazid, rifampicin, ethambutol, pyrazinamide, and streptomycin; three injectable second-line drugs— amikacin, capreomycin, and kanamycin; three fluoroquinolones— ciprofloxacin, moxifloxacin, and ofloxacin; and one less commonly used second-line drug, ethionamide (Table 1) [7, 26]. The short reads containing the whole-genome sequences of these 7,845 isolates were downloaded from the European Nucleotide Archive [21] and the Sequence Read Archive [22]. The accession numbers which we used in our dataset were ERP[000192, 006989, 288008667, 010209, 013054, 000520], PRJEB[10385, 10950, 14199, 2358, 2794, 5162, 9680], PRJNA[183624, 235615, 296471], and SRP[018402, 051584, 061066]. This is the same dataset we used in a previous study, whose focus was on the development of interpretable drug resistance prediction models [45].

**Table 1:** The number of isolates and the label distribution in our data.

| Drug | Num of labelled isolates | Num of resistant isolates |
| --- | --- | --- |
| Streptomycin | 5,125 | 2,104 (41%) |
| Rifampicin | 7,715 | 2,968 (38%) |
| Pyrazinamide | 3,858 | 754 (20%) |
| Ofloxacin | 2,911 | 800 (27%) |
| Moxifloxacin | 961 | 129 (13%) |
| Kanamycin | 2,436 | 697 (27%) |
| Isoniazid | 7,734 | 3,445 (45%) |
| Ethionamide | 1,516 | 498 (33%) |
| Ethambutol | 6,096 | 1,407 (23%) |
| Ciprofloxacin | 443 | 37 (8%) |
| Capreomycin | 1,991 | 552 (28%) |
| Amikacin | 2,033 | 573 (28%) |

We used an additional, independent, dataset to test how well the results generalize. We downloaded the SNP-based feature matrix of a dataset from the British Columbia Centre for Disease Control (BC-CDC), used in another publication by our group [14]; we describe this dataset in more detail in Table 2. This dataset contains over 1,138 TB isolates with their labels for the five first-line drugs: isoniazid, rifampicin, ethambutol, pyrazinamide, and streptomycin. We used this dataset for the results shown in Figure 4, and the PATRIC and ReSeqTB datasets for the remainder of the experiments.

**Figure 1:**
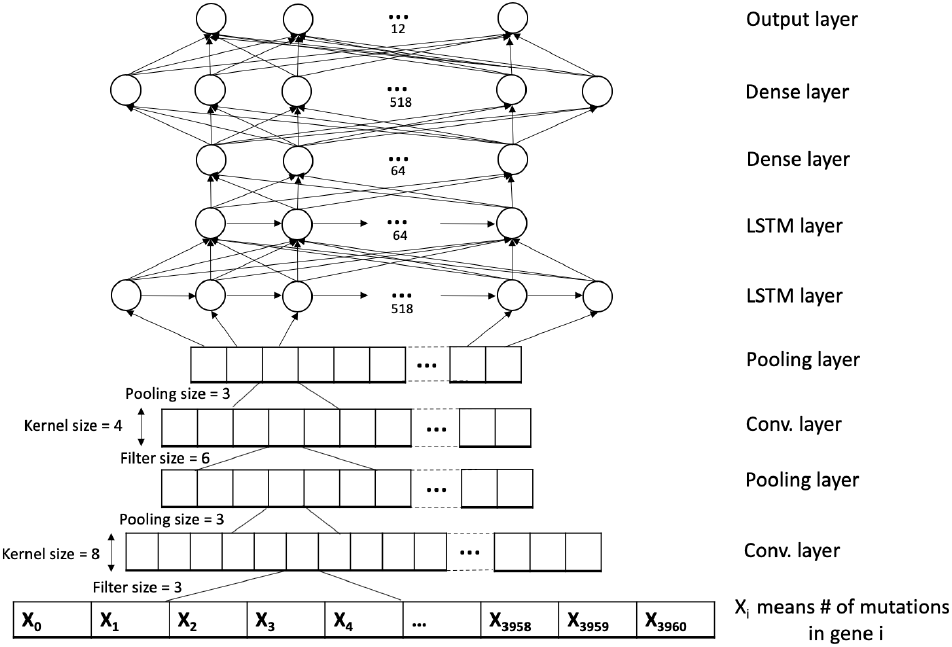
The architecture of our LRCN network.

**Figure 2:**
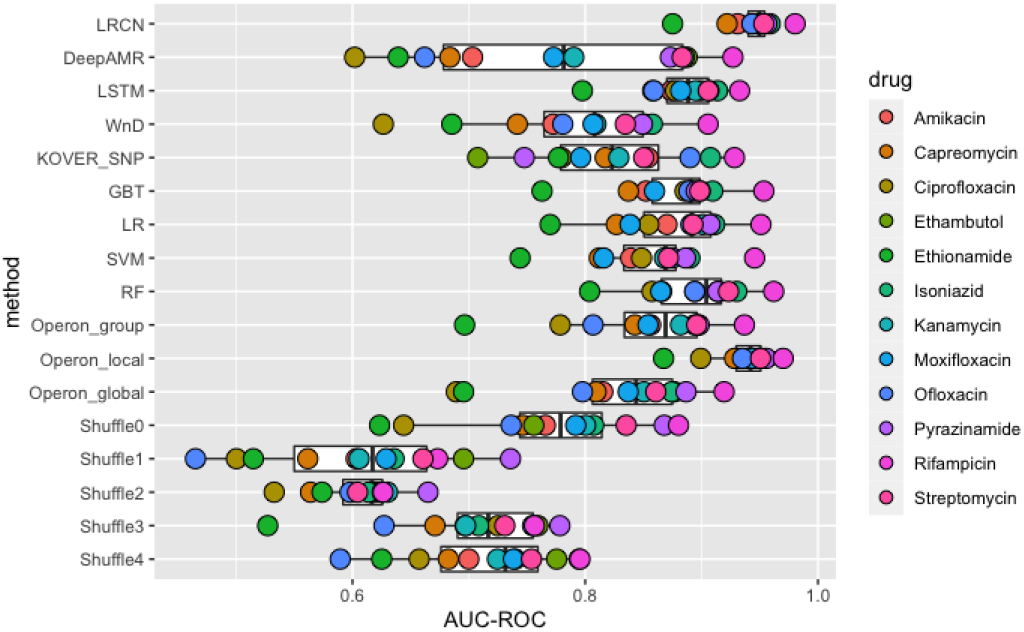
Performance of all the methods we tested, nonnested cross-validation approach. The vertical axis lists the methods used, each dot represents a drug, and the horizontal axis shows the AUC-ROC. The white rectangles represent the mean and standard deviation of each method. The “Operon” and “Shuffle” methods are described in Sections 3.2 and 3.4, respectively.

**Figure 3:**
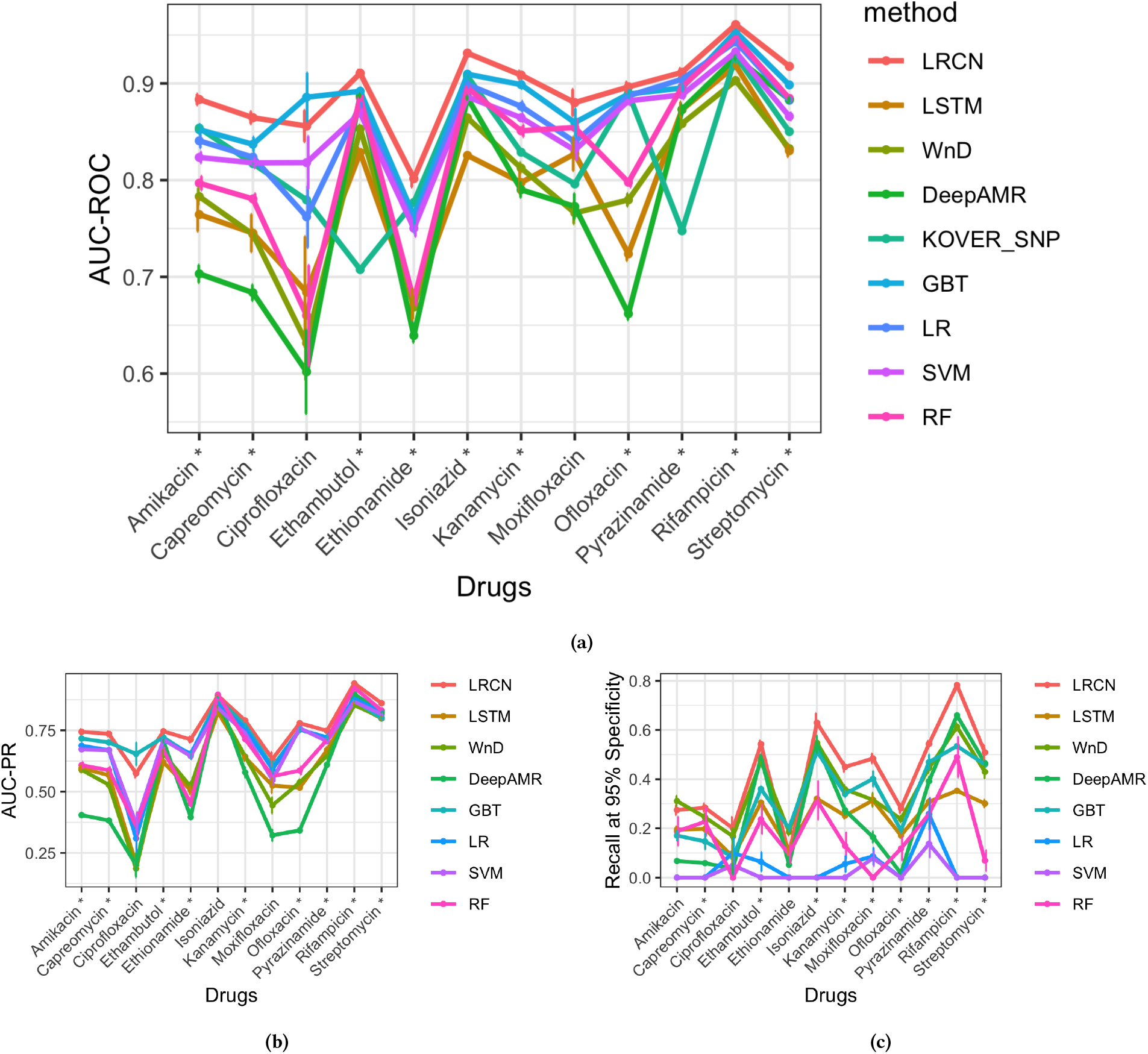
Comparison of LRCN to state-of-the-art models. The error bars represent the confidence intervals on the performance, calculated using a nested cross-validation approach (Section 2.4). We calculated 95% confidence interval as the mean ±1.96 times the standard error across the 10 performance values. Statistically significant differences between LRCN and the second-best method are marked with a “*”. See Supplementary Table 12 for more details of these results. The vertical axis indicates: (a) AUC-ROC, (b) AUC-PR and (c) sensitivity at 95% specificity.

**Figure 4:**
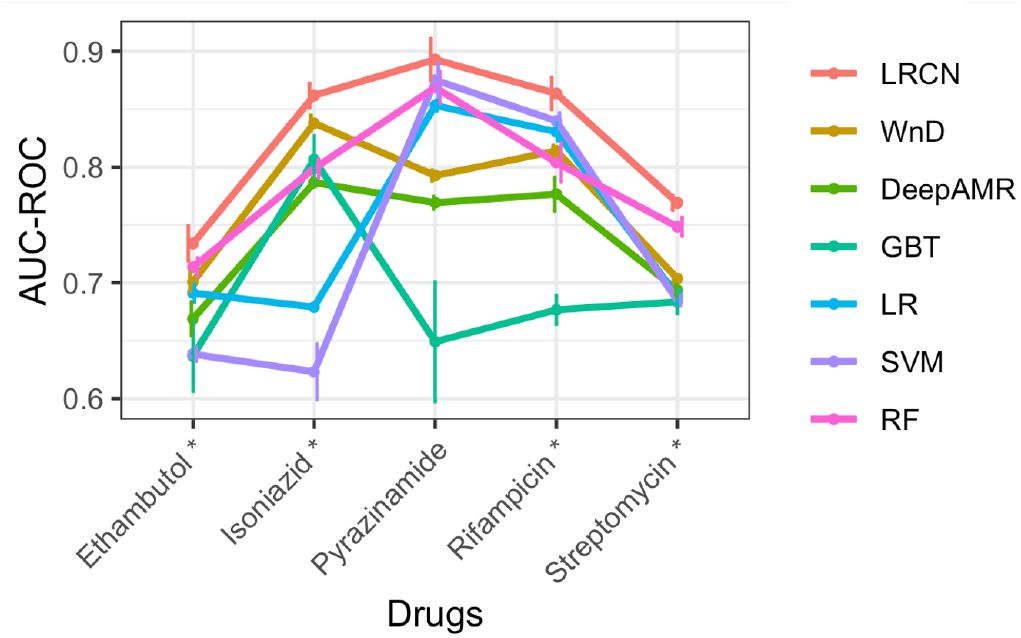
Comparison of LRCN to state-of-the-art models on the BCCDC data. Statistically significant differences are marked with a “*”.

**Table 2:** Summary of the number of isolates and the label distribution in BCCDC data.

| Drug | Num of labelled isolates | Num of resistant isolates |
| --- | --- | --- |
| Streptomycin | 1,136 | 87 (7.65%) |
| Rifampicin | 1,138 | 26 (2.28%) |
| Pyrazinamide | 134 | 13 (9.7%) |
| Isoniazid | 1,138 | 157 (13.79%) |
| Ethambutol | 1,138 | 22 (1.96%) |

### 2.2 Variant calling

To obtain the SNP information from the raw reads of the PATRIC and ReSeqTB datasets, we used a standard protocol [5, 7]. We first mapped the raw sequence data to the reference genome, *Mycobacterium tuberculosis H37Rv*, using bwa-mem [23], then called the variants (SNPs, insertions, and deletions) for each isolate using two different established pipelines, SAMtools [24] and GATK [31]. To make the calls more robust, we used the intersection of variant calls between the two tools; fewer than 4% of the variants were removed in this step. These SNP-based features were then represented as a binary matrix of 7,845 isolates (in rows) by 742,620 variants (in columns). The BCCDC dataset was processed into variant calls [14] before we acquired it, so we skipped this step for that dataset.

We created gene burden features by identifying, for a given isolate, the number of variants in each gene. We acquired boundaries for each known TB gene from MycoBrowser [18]. Of the 4,187 known TB genes, we found that 3,960 had a variant in at least one isolate in our dataset. We represent these gene burdens of thePredicting drug resistance in *M. tuberculosis* using a Long-term Recurrent Convolutional Network PATRIC and ReSeqTB datasets as an integer 7,485 by 3,960 matrix, in which each entry indicates the number of mutations in a given gene within a given isolate.

The advantage of using gene burden features instead of SNP-based features is that gene burden data drastically reduces the number of features, mitigating the “curse of dimensionality”. When the number of available isolates is small relative to the number of SNPs, using the gene burden data may lead to more accurate models and reduce overfitting. The gene burden-based method may lose this advantage in situations when much larger sample sizes are available.

### 2.3 LRCN model

Our method is based on a combination of long short-term memory (LSTM) [15] layers and convolutional neural network (CNN) [34] layers, an architecture known as a Long-term Recurrent Convolutional Network (LRCN) [9]. An LSTM is an artificial recurrent neural network that can learn long-distance dependencies by using feedback connections, which is appropriate for our situation given the multifactorial nature of the drug resistance phenotype [39]. More precisely, it is known that drug resistance in *M. tuberculosis* is mediated by compensatory mutations, which help reduce the fitness cost of drug-resistance causing mutations, but may occur in a different gene.

A CNN is a feed-forward neural network designed for processing structured arrays of data, especially data whose linear order is important. We elaborate on this further in Section 3.2, where we show that the linear order is important for our gene burden-based features. The combination of a CNN with an LSTM enables the model to take into account both the linear order as well as the local structure of the genes in a genome.

The architecture of our model is as follows. The input for each isolate is encoded as an integer vector *I* = {*g*_1_,…, *g_m_*}, where *m* is the number of genes used (here, *m* = 3, 960) and *g_i_* is the number of SNPs the isolate has in the *i*-th gene. The output for this isolate is a binary vector *O* = {*d*_1_,…, *d_n_*} where *n* is the number of drugs (here, *n* = 12), and *d_j_* is the isolate’s predicted status for the *j*-th drug. This input is first processed by *L_conno_* = 2 convolutional layers (`keras.layers.Conv1D` and `keras.layers.MaxPooling1D`) with specific filter (*f_i_*), kernel (*k_i_*), and pool (*p_i_*) sizes, where *i* represents the layer number. These layers use the rectified linear unit (ReLU) activation function and the “same” padding (padding evenly to the left/right of the input such that the output has the same width dimension as the input). The output of the last CNN layer is processed by *L_lstm_* = 2 LSTM layers (`keras.layers.LSTM`), with a specified number of nodes (*lstm_i_*). They use the hyperbolic tangent activation function and a recurrent dropout with a rate of *p_rec_* = 0.3. At the end, the output of the LSTM feeds into the dense layers (`keras.layers.Dense` with a specified number of nodes (*dense_i_*), and the last layer generates the output vector. Between every pair of layers, we use a dropout (`keras.layers.Dropout`) with a rate of *p* = 0.1 to reduce overfitting. A schematic representation of this architecture is shown in Figure 1.

The parameters of the optimized model (for the non-nested crossvalidation approach in section 2.4) are as follows. For the CNN layers, the parameters are (*f*_1_, *k*_1_, *p*_1_) = (8, 3,3) and (*f*_2_, *k*_2_, *p*_2_) = (4, 6,4). The LSTM layers have *lstm*_1_ = 518 and *lstm*_2_ = 64 nodes, respectively. Finally, the dense layers have *dense*_1_ = 64 and *dense*_2_ = 518 nodes, respectively, and the last dense layer has *n* = 12 nodes to generate the output vector (Figure 1). We use the Adam optimizer with a learning rate of *η* = 0.01 to train the model. The parameters chosen by the nested cross-validation approach (Section 2.4) are listed in Supplementary Table 4.

We use a multi-task model, which predicts an isolate’s resistance status for all 12 drugs in a single network. This approach’s advantage is that if two drugs have a shared structure, either through highly correlated resistance status vectors due to their joint use in clinical regimens or through common resistance mechanisms, then the patterns learned from one drug can compensate for a smaller amount of training data for the other drug. The pairwise correlation of the status vectors for each pair of drugs is shown in the Supplementary Section 2.

The challenge of using the multi-task model in our dataset is that many isolates lack labels for some of the drugs. For this reason we replace the usual loss function with a masked loss function. The masked loss function ignores the missing labels in calculating the loss, and therefore, these missing labels do not affect the network weights. We use the binary cross-entropy as the loss function. Specifically, we use the loss function

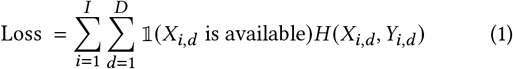

where *I* and *D* are the numbers of isolates and drugs respectively, *X_i,d_* and *Y_i,d_* are the true and predicted resistance values, *H* is the binary cross-entropy function, and 1 is the indicator function.

### 2.4 Train-Validation-Test split

To evaluate our method, we split our data into training, validation, and test sets. We use two different approaches for this purpose, as we detail below.

To obtain the results shown in Figure 3 and 4, we used a nested cross-validation approach. We split the data into 10 equal parts, and trained the model 10 times, using 8 folds for training, 1 fold for validation, and 1 for testing each time. In each run, we used Bayesian Optimization (see Section 2.6) to maximize the Area Under The Receiver Operating Characteristic Curve (AUC-ROC) on the validation set, and after tuning the hyperparameters, we tested the best model on the test set. At the end of this process, we ended up with 10 different models for each method, from which we determined the mean and standard error of the AUC-ROC.

Because the approach above is computationally expensive, we used regular (non-nested) cross-validation for figures other than Figure 3 and 4. We used 10% of the data as the testing set. We chose hyperparameters via 10-fold cross-validation on the remaining dataset by selecting the model that achieved the highest mean AUC-ROC. After hyperparameter tuning, we evaluated our model on the test set.

In both approaches, we used stratified *k*-fold cross-validation so that each fold had an equal fraction of resistant and susceptible samples.

### 2.5 Evaluation

In order to evaluate the accuracy of our predictions, we used the AUC-ROC and AUC-PR values. The AUC-ROC metric is the area under the plot of the true positive rate (TPR) against the false positive rate (FPR) at different classification thresholds. The AUC-PR is similar to AUC-ROC, but the plot is that of precision 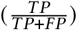 against recall 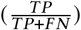 at different classification thresholds.

Since many clinical applications require a predictor with at least a 95% specificity [38], we also evaluate our method using the sensitivity at 95% specificity. This metric tells us what fraction of resistant isolates can be identified as such when that their threshold for prediction is set so as to give at least 95% specificity. Several methods do not achieve a 95% specificity at any threshold, meaning that they cannot be used when a 95% specificity is required.

### 2.6 Hyperparameter optimization

For tuning the model hyperparameters and determining the ideal number of layers for each type of subnetwork (CNN, LSTM, dense layer) in our model we used Bayesian optimization. Bayesian optimization is a method suited to optimizing high-cost functions, such as hyperparameter search for deep learning models, by using a combination of a Gaussian process and Bayesian inference [35]. We used the Python implementation of Bayesian Optimization [27] with 15 iterations to tune the hyperparameters of all the models used in this paper, with a uniform optimization approach to ensure a fair comparison.

### 2.7 Existing methods

We compared our LRCN model with a large number of existing methods that display state-of-the-art performance. We omitted catalog methods and other methods that have been found to perform significantly worse than the methods listed here. The following methods were used for the comparison:

- DeepAMR [43]. This method is optimized for predicting drug resistance in TB.
- WnD (Wide-n-Deep neural network) [3]: This method is optimized for predicting drug resistance in TB.
- KOVER (Set Covering Machine algorithm) [10]: A rule-based model to predict drug resistance in multiple bacterial species, including TB.
- GBTs (Gradient-Boosted Trees) [7, 20]: Some publications suggest that this method has the best performance in drug resistance studies, especially in TB.
- LR (Logistic regression) [20]: Some recent research demonstrates their superior performance in predicting drug resistance in TB.
- RF (Random Forests) [19]: Based on some prior research, RFs can achieve very good performance in predicting drug resistance in TB.
- SVM (Support Vector Machines): SVMs are a widely used ML model for binary classification, with applications in many areas.
- LSTM (Long Short-Term Memory) is a widely used neural network model. Note that our LRCN model includes an LSTM component.

Since the publications that propose these models used SNP-based features, we evaluated all the methods separately using SNP-based features and separately using gene burden-based features, to compare the relative performance of these two sets of features with the same class of models in addition to comparing the models to each other (Section 3.3).

For all the comparison models we used the exact same data and procedure (e.g. Bayesian Optimization to choose the parameters) that we used for the LRCN. The parameters chosen for the nested cross-validation approach (Section 2.4) are listed in Supplementary Section 3. Because KOVER produces discrete classifications and does not output a score, its AUC-ROC is derived from an ROC curve with a single point. Also, it is not possible to calculate the AUC-PR and the sensitivity at 95% specificity with KOVER’s output (see Supplementary Section 3.1).

The optimized parameters for the non-nested cross-validation were as follows:

- WnD: 5 layers with 518, 518, 64, 518, and 64 nodes, respectively, and a kernel regularizer value of 0.1.
- GBT: 30 estimators, *min_samples_split* = 4, *maxdepth* = 130, and *random_state* = 1
- LR: the *ℓ*_2_ penalty and *C* = 0.1.
- RF: 140 estimators, a minimum sample split of 4, no bootstrapping, and a maximum depth of 50
- SVM: a linear kernel and *C* = 0.1
- LSTM: 3 LSTM layers, with 355, 455, and 343 nodes, respectively, followed by 4 dense layers with 359, 219, 230, and 147 nodes, respectively.

The parameters chosen by the nested cross-validation approach (Section 2.4) are listed in Supplementary Section 3.

### 2.8 Implementation

We used the Python programming language to implement all the methods in this paper. We used the Keras library [4] for the deep neural networks, and the Scikit-learn library [30] for the machine learning models.

## 3 RESULTS

Figure 2 summarizes the performance of most of our models, its various permutations, and the models we compared it with. In this section we describe each comparison in more detail.

### 3.1 LRCN outperforms state-of-the-art methods on a large dataset

We evaluated our LRCN method by comparing it to other existing methods, namely, DeepAMR, WnD, KOVER (which uses the SCM algorithm), LSTM, GBT, RF, LR, and SVM (Section 2.7). In our evaluation we investigated the following questions:

- Does gene burden-based LRCN outperform other existing methods on the AUC-ROC and AUC-PR metrics? If so, is the difference statistically significant?
- Does a gene burden-based LRCN architecture generalize across multiple data sets?
- How useful is the gene burden-based LRCN for clinical applications? We evaluate this by using a novel metric, the sensitivity at a 95% specificity.

We found that the LRCN method achieves a substantial improvement over all existing methods in most cases. For the AUC-ROC metric, we found that the LRCN method achieves better performance than all the state-of-the-art methods for 10 of the 12 drugs (Figure 3a). For the AUC-PR metric, the LRCN method achieves better performance for 9 of the 12 drugs (Figure 3b). This performance improvement is statistically significant; a one-sided *t*-test achieves a *p*-value below 0.05 (see Supplementary Table 11). The difference in performance between LRCN and other methods on moxifloxacin in both AUC-ROC and AUC-PR, ciprofloxacin in AUC-ROC, and isoniazid in the AUC-PR metric is not statistically significant, so we cannot say whether LRCN performs better or worse than other methods. However, for ciprofloxacin, GBT performs statistically significantly better in AUC-PR. This poor result on ciprofloxacin may be due to the limited data available for that drug, for which we only have 37 resistant samples.

To gain further confidence in LRCN’s robustness and to make sure that our model generalizes to other datasets, we additionally evaluated the trained models on the BCCDC dataset. This data set differs from ReSeqTB and PATRIC in its geographical homogeneity, as it represents a single province in Canada. We observed that the gene burden-based LRCN performed statistically significantly better on 4 of 5 drugs (Figure 4). The difference in performance between LRCN and other methods on pyrazinamide is not statistically significant, which may be because only 134 samples in the BCCDC dataset have a status available for pyrazinamide.

For a prediction of resistance to be usable for clinical diagnosis, it must be made with a high specificity, especially for first-line drugs because the second-line drugs tend to cause more severe side effects and may lead to more frequent treatment failure [11, 46]. Therefore, we also evaluated methods according to the maximum sensitivity they achieve with a minimum of 95% specificity (Section 2.5). That is, we find the smallest threshold at which the model has 95% specificity, then we calculate the sensitivity at this threshold. If the model is unable to reach 95% specificity for any threshold, it cannot be used for diagnosis, and therefore scores zero in this metric. The LRCN method has the best sensitivity among all the methods on 9 of the 12 drugs and this difference in performance is statistically significant (*p* < 0.05, one-sided *t*-test; see Supplementary Table 11). Although for ciprofloxacin, LRCN’s performance difference relative to other methods is not statistically significant, and for the two remaining drugs, amikacin and ethionamide, the WnD method achieves a slightly higher sensitivity. Furthermore, many standard ML approaches perform poorly on this metric, which may limit their usefulness in clinical applications (Figure 3c).

To evaluate whether the differences in model performance are due to random chance, we calculated the *p*-values as well as the 95% confidence intervals for each of our three metrics: AUC-ROC, AUC-PR, and sensitivity at 95% specificity. Since we used 10-fold cross-validation, we came up with 10 different model performance metrics for each method (Section 2.4). We calculated 95% confidence interval as the mean ±1.96 times the standard error across the 10 performance values. We calculated the *p*-values using a one-sided *t*-test between the vectors of size 10 holding these performance values for our method and the competitor methods.

### 3.2 LRCN exploits gene order

We hypothesized that the improvement in performance in LRCN is due to the importance of gene order. Indeed, the main difference between the LRCN and the other methods is the combination of CNN and LSTM layers. These layers may help the LRCN to recognize the genes’ spatial relationships in the genome.

To test this hypothesis, we randomly permuted the genes to put them in a random order. We then trained and tested our method with the new data. This permutation is analogous to shuffling the pixels of an image. We shuffled the genes’ order five times and for each new permutation, we carried out the entire process including Bayesian Optimization hyperparameter tuning.

As expected, we found a significant decrease in the LRCN’s performance on the shuffled data, making it comparable to that of a random model. This implies the importance of genes’ order in our model (Figure 5).

**Figure 5:**
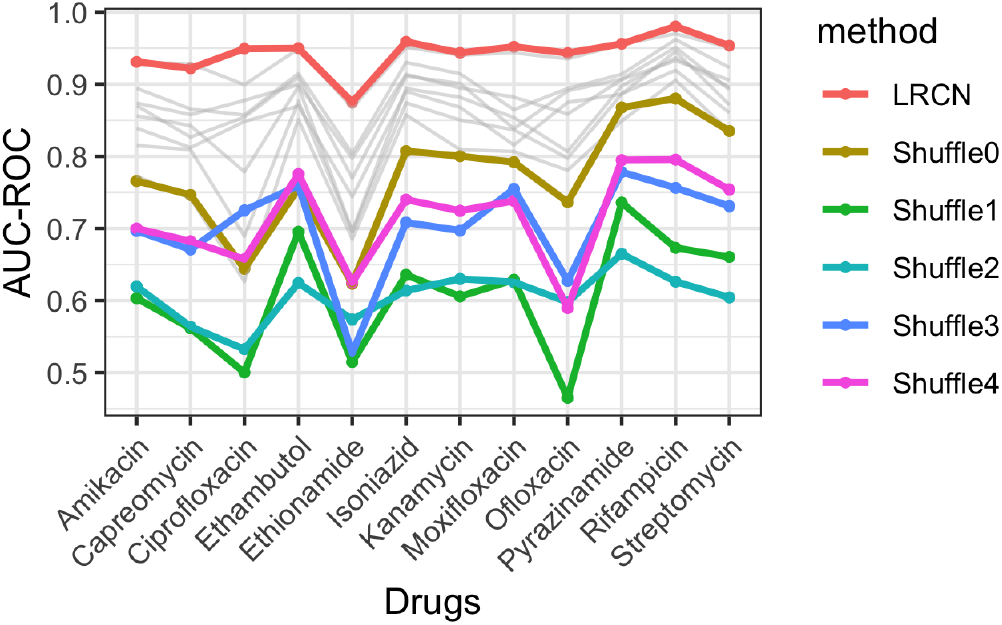
Comparison between the test AUC-ROC of our LRCN model using regular and shuffled gene orders. The gray lines represent the result of other models.

### 3.3 Gene burden-based features enable a high accuracy on our dataset

To evaluate the efficacy of the gene burden-based approach, we applied the same methods to the SNP-based features (Section 2.7). Using SNP-based features, we trained and tested the LRCN and all state-of-the-art methods [3, 7, 10, 19, 20, 43]. We found that the LRCN approach on the SNP-based features performed similarly to other methods (Figure 6b). However, all SNP-based approaches underperform the gene burden-based LRCN (Figure 6a). This suggests that using either just gene burden features or just LRCN does not achieve the best performance; both techniques together are required for improved performance.

**Figure 6.**
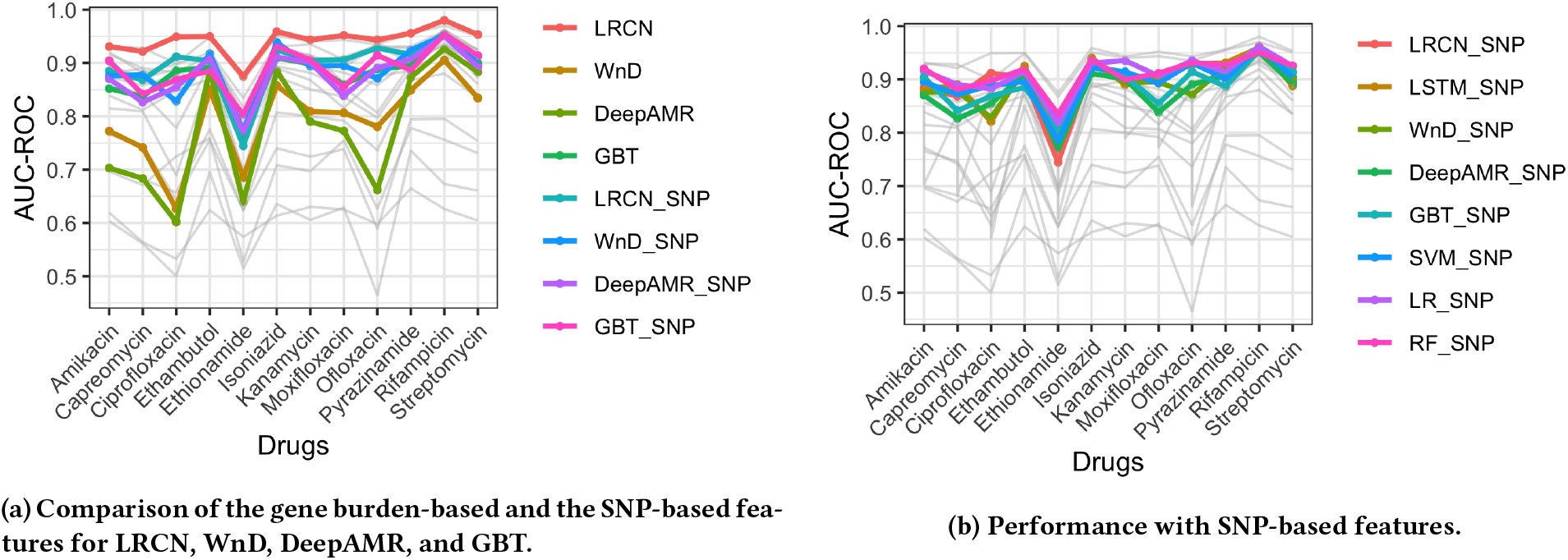

### 3.4 The organization of genes into operons drives performance

We hypothesized that the LRCN leverages the fact that neighboring genes in operons have related functions. Operons are clusters of neighbors or co-regulated genes with related functions [29]. Among the 3960 genes with at least one mutation in our data, 879 genes fall within an operon. For this experiment, we shuffled the operon genes in three different ways to disrupt their order. For each one, we trained and tested our method on the resulting shuffled (permuted) data.

First, we hypothesized that the order of an operon’s genes has a greater effect on the LRCN’s results comparing to the order of non-operon genes. To test this, we randomly permuted all the genes found within operons while leaving the other genes intact. We found a significant decrease in the accuracy of the LRCN, similar to that seen when we randomly permute all the genes (Figure 7). This suggests the importance of gene order within operons in comparison to that of non-operon genes.

**Figure 7:**
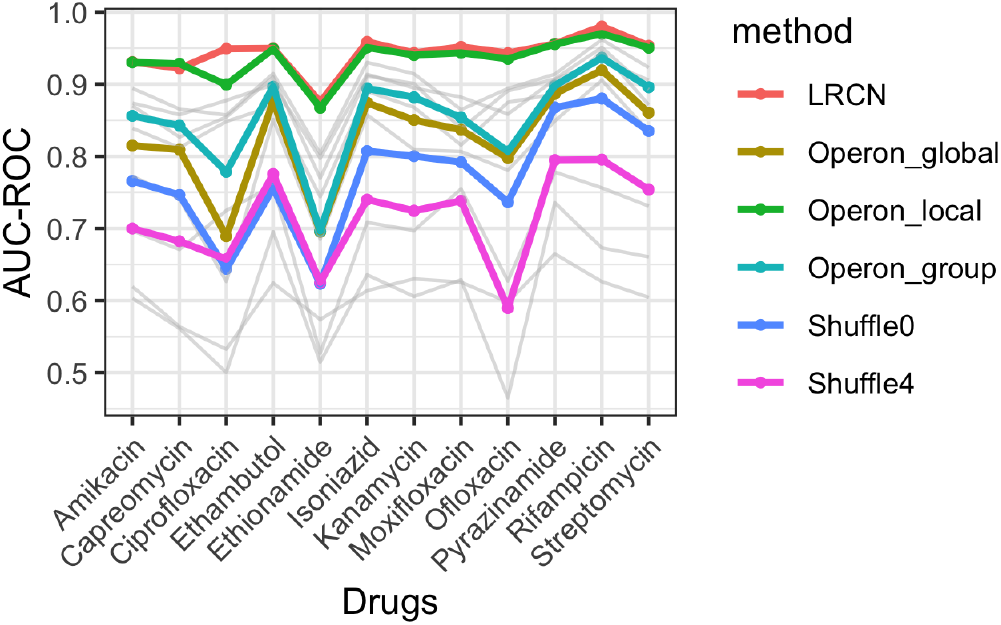
Evaluation of LRCN performance on permuted data sets. “operon_global” is the first approach described in Section 3.4, “operon_local” is the second approach, and “operon_group” is the third approach. “Shuffle0” and the “Shuffle4” are the highest accuracies from section 3.2.

Second, we assumed that as the operons are typically short (from 2 to 14 genes), the order of genes within each operon should not affect the model’s performance, and we should not expect a considerable difference with the original result. To test this, we individually permuted the genes within each operon, while leaving all the other genes untouched. We observed that, as hypothesized, with this approach the LRCN’s performance is comparable to its performance on the original data (Figure 7).

Our third hypothesis was that if we keep the gene order fixed within each operon, but permute the locations of the operons, this should negatively affect the accuracy of the LRCN (this is analogous to shuffling the frames of a video, which could render the videos meaningless). We have indeed found a considerable drop in the LRCN’s performance in this case (Figure 7).

### 3.5 Sharing information between drugs improves performance

We hypothesized in section 2 that using a multi-task model could improve the accuracy of our model, especially for drugs with relatively limited labeled data available, driven by shared mechanisms of resistance. If true, this hypothesis would imply that the patterns learned from one drug can compensate for the lack of training data for another drug. We observed that using the multi-task model improved the performance of the gene burden-based LRCN (Figure 8). Furthermore, we see that the gene burden-based LRCN trained on a single drug still displays good performance and can accurately predict drug resistance to that specific drug.

**Figure 8:**
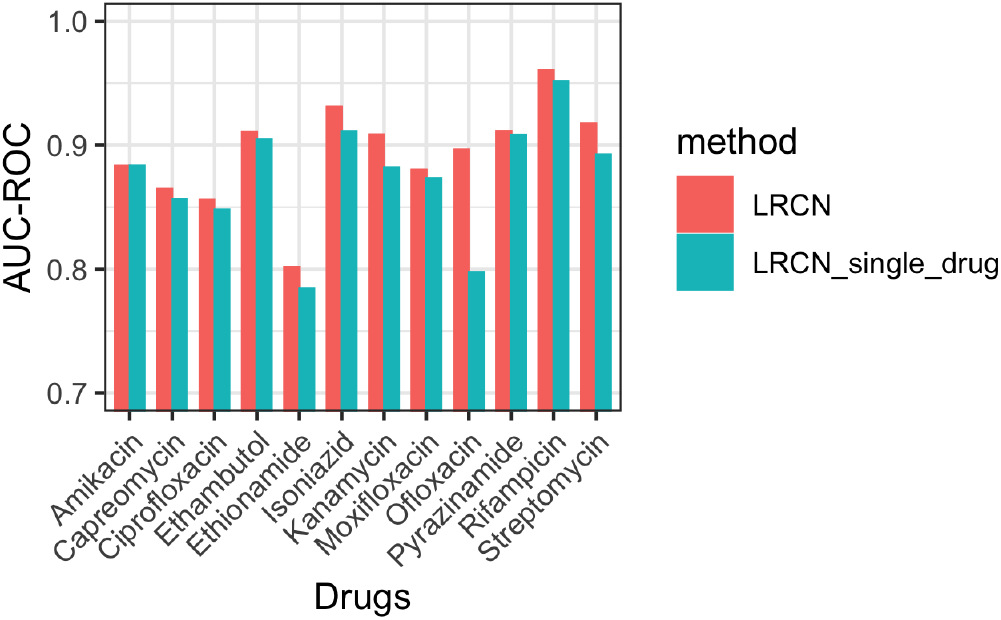
Comparison of the performance of the multi-task LRCN and the single-drug LRCN, using gene burden-based features.

## 4 CONCLUSION

In this paper, we propose the novel use of a deep learning architecture, LRCN, for predicting drug resistance in *M. tuberculosis*. This architecture is a combination of CNN and LSTM layers and thus has the advantage of considering both the gene mutation burden and the gene order.

Our results suggest three major findings:

- Gene burden-based LRCN outperforms the other existing methods in a variety of settings, using three different evaluation metrics, including one relevant for clinical applications. This good performance of the gene burden-based LRCN is robust, as evidenced by a variety of additional experiments including a test on an independent dataset.
- We observed that LRCN achieves its highest performance when we use the gene burden, defined as the number of mutations in each gene, rather than the full SNP data. The gene burden allows us to take into account the importance of individual genes, while the model architecture allows us to consider their order in the genome.
- We found that using the CNN layers before the LSTM layers in our architecture is beneficial for drug resistance prediction, which illustrates the importance of gene locality. We established the importance of gene order with two different experiments. Importantly, these experiments suggest that the good performance of the LRCN model on TB isolates is robust and not explained by chance alone.

In conclusion, we introduced a novel state-of-the-art method for predicting drug resistance by using gene order information alongside gene burden information. Our results suggest that gene burden-based prediction is effective in the context of an LRCN.

This study also suffers from several limitations that we hope to address in future work, namely:

- This paper focuses exclusively on *M. tuberculosis* as a proof of concept for the approach. In future work, we plan to apply the LRCN approach to predict the drug resistance in other pathogens such as *Escherichia coli*.
- In this paper, the focus was on achieving the most accurate model for predicting drug resistance. However, the lack of interpretability is a considerable disadvantage of the LRCN model as well as most other neural network-based approaches. For this reason, we plan to add information to the LRCN method to make it more interpretable in future work.

## 5 ACKNOWLEDGEMENTS

Leonid Chindelevitch acknowledges funding from the MRC Centre for Global Infectious Disease Analysis (reference MR/R015600/1), jointly funded by the UK Medical Research Council (MRC) and the UK Foreign, Commonwealth & Development Office (FCDO), under the MRC/FCDO Concordat agreement and is also part of the EDCTP2 programme supported by the European Union. This project was funded by the Genome Canada grant 283BAC, “Machine learning methods to predict drug resistance in pathogenic bacteria”.

## Footnotes

1 https://github.com/AmirHoseinSafari/Genotype-collector-and-SNP-dataset-creator

2 https://github.com/AmirHoseinSafari/LRCN-drug-resistance

3 https://github.com/AmirHoseinSafari/M.tuberculosis-dataset-for-drug-resistant

4 https://github.com/AmirHoseinSafari/Predicting-drug-resistance-in-M.-tuberculosis-using-a-Long-term-Recurrent-Convolutional-Network/blob/main/Supplementary-Materials.pdf

