## Supplementary-Materials for "Predicting drug resistance in *M. tuberculosis* using a Long-term Recurrent Convolutional Network"

### Supplementary Material of Predicting drug resistance in *M. tuberculosis* using a Long-term Recurrent Convolutional Networks architecture

#### 1 Related works

Title, Author, Short description

1. Predicting antimicrobial resistance using conserved genes, Marcus Nguyen, AMR via conserved genes
2. Genomic prediction of tuberculosis drugresistance: benchmarking existing databases and prediction algorithms, Tra-My Ngo, Benchmarking TB databases
3. DeepAMR for predicting co-occurrent resistance of Mycobacterium tuberculosis, Yang Yang, DeepAMR paper
4. Genetic Determinants of Drug Resistance in Mycobacterium tuberculosis and Their Diagnostic Value, Maha R. Farhat, DecisionTreesForTB.
5. Deciphering drug resistance in Mycobacterium tuberculosis using wholegenome sequencing: progress, promise, and challenges, Keira A. Cohen, Deciphering drug resistance in Mycobacterium tuberculosis
6. Prediction of Susceptibility to First-Line Tuberculosis Drugs by DNA Sequencing, , CRyPTiC susceptibility predictions
7. Combining Structure and Genomics to Understand Antimicrobial Resistance, Tanushree Tunstall, Combining Structure and Genomics to Understand Antimicrobial Resistance
8. Beyond multidrug resistance: Leveraging rare variants with machine and statistical learning models in Mycobacterium tuberculosis resistance prediction, Michael L. Chen,
9. Prediction of antibiotic resistance in Escherichia coli from large-scale pan-genome data, Danesh Moradigaravand, E coli resistance prediction
10. Prediction of Acquired Antimicrobial Resistance for Multiple Bacterial Species Using Neural Networks, Aytan-Aktug, Drug resistance via DNN
11. Evaluation of Whole-Genome Sequence Method to Diagnose Resistance of 13 Anti-tuberculosis Drugs and Characterize Resistance Genes in Clinical Multi-Drug Resistance Mycobacterium tuberculosis Isolates From China, Xinchang Chen, Drug resistance prediction tools

12. Deciphering drug resistance in Mycobacterium tuberculosis using wholegenome sequencing: progress, promise, and challenges, Keira A. Cohen, DR-TB via WGS

13. Prediction of rifampicin resistance beyond the RRDR using structure-based machine learning approaches, Stephanie Portelli, DR via structure-based machine learning

14. Rapid inference of antibiotic resistance and susceptibility by genomic neighbour typing, Karel Břinda , DR by genomic neighbour typing

15. Detection of low-frequency resistance-mediating SNPs in next-generation sequencing data of Mycobacterium tuberculosis complex strains with binoSN, Viola Dreyer, Detection of low-level TB resistance

16. Interpreting k-mer–based signatures for antibiotic resistance prediction, Magali Jaillard, K-mer interpretation

17. Interpretable Model for Antibiotic Resistance Prediction In Bacteria Using Deep Learning, MANOJ JHA, Interpretable Model for Antibiotic Resistance Prediction In Bacteria Using Deep Learning

18. Improved Resistance Prediction in Mycobacterium tuberculosis by Better Handling of Insertions and Deletions, Premature Stop Codons, and Filtering of Non-informative Sites, Camilla Hundahl Johnsen, Improved Resistance Prediction in Mycobacterium tuberculosis

19. GWAS for quantitative resistance phenotypes in Mycobacterium tuberculosis reveals resistance genes and regulatory regions, Maha R. Farhat, GWAS for quantitative resistance phenotypes in Mycobacterium tuberculosis reveals resistance genes

20. Genetics and roadblocks of drug resistant tuberculosis, João Perdigão, Genetics-and-roadblocks-of-drug-resistant-tu-2019-Infection–Genetics-and-Ev

21. The role of whole genome sequencing in antimicrobial susceptibility testing of bacteria: report from the EUCAST Subcommittee, M.J. Ellington, EUCAST report

22. Rapid genomic first- and second-line drug resistance prediction from clinical Mycobacterium tuberculosis specimens using Deeplex®-MycTB, Silke Feuerriegel, ERJ paper on rapid DR prediction

23. pncA gene mutations in reporting pyrazinamide resistance among the MDRTB suspects, Xiaoyuan Wu, pncA-gene-mutations-in-reporting-pyrazinamide-resi-2019-Infection–Genetics

24. Next-Generation Sequencing Approaches to Predicting Antimicrobial Susceptibility Testing Results, Rebecca Yee, NGS for DR prediction

25. WGS to predict antibiotic MICs for Neisseria gonorrhoeae, David W. Eyre, Neisseria dataset with MICs

26. Using machine learning to predict antimicrobial minimum inhibitory concentrations and associated genomic features for nontyphoidal Salmonella, Marcus Nguyen, MIC prediction in Salmonella

27. Multi-Label Random Forest Model for Tuberculosis Drug Resistance Classification and Mutation Ranking, Samaneh Kouchaki ,

28. A large scale evaluation of TBProfiler and Mykrobe for antibiotic resistance prediction in Mycobacterium tuberculosis, Pierre Mahé, Mahe Tournoud comparison of TB tools

29. Application of machine learning techniques to tuberculosis drug resistance analysis, Samaneh Kouchaki, LogisticRegressonForTB

30. Large scale modeling of antimicrobial resistance with interpretable classifiers, Alexandre Drouin, Kover latest paper

31. Interpretable genotype-tophenotype classifers with performance guarantees, Alexandre Drouin, Kover paper

32. Developing an in silico minimum inhibitory concentration panel test for Klebsiella pneumoniae, Marcus Nguyen, Klebsiella MICs

33. Robust detection of point mutations involved in multidrug-resistant Mycobacterium tuberculosis in the presence of co-occurrent resistance markers, Julian Libiseller-Egger, Robust detection of point mutations involved in MDR-TB

34. Systematic review of mutations associated with resistance to the new and repurposed Mycobacterium tuberculosis drugs bedaquiline, clofazimine, linezolid, delamanid and pretomanid, Suha Kadura, Resistance to new TB drugs

35. Machine learning with random subspace ensembles identifies antimicrobial resistance determinants from pan-genomes of three pathogens, Jason C. HyunI, Random subspace ensembles for AMR

36. Recent advances in molecular diagnostics and understanding mechanisms of drug resistance in nontuberculous mycobacterial diseases, Hee Jae Huh, Recent-advances-in-molecular-diagnostics-and-understandi-2019-Infection–Gen

37. Prediction of pyrazinamide resistance in Mycobacterium tuberculosis using structure-based machine learning approaches., Joshua J Carter, Pyrazinamide resistance in TB

38. Predicting antimicrobial resistance in Pseudomonas aeruginosa with machine learning-enabled molecular diagnostics, Ariane Khaledi, Pseudomonas genomics + transcriptomics of DR

39. Predicting Phenotypic Polymyxin Resistance in Klebsiella pneumoniae through Machine Learning Analysis of Genomic Data, Nenad Macesic, Predicting resistance in Klebsiella

40. Whole-Genome Sequencing for Drug Resistance Profile Prediction in Mycobacterium tuberculosis, Sebastian M. Gygli, WGS for MTB DR

41. Evaluation of parameters affecting performance and reliability of machine learning-based antibiotic susceptibility testing from whole genome sequencing data, Allison L. Hicks,

42. Structure guided prediction of Pyrazinamide resistance mutations in pncA, Malancha Karmakar, Structure-guided PZA resistance prediction

43. Survey of drug resistance associated gene mutations in Mycobacterium tuberculosis, ESKAPE and other bacterial species,AbhirupaGhosh,SurveyOfMutations

44. Machine Learning Predicts Accurately Mycobacterium tuberculosis Drug Resistance From Whole Genome Sequencing Data, Wouter Deelder, Taane Clark's latest paper on ML for DR

45. Prediction of Phenotypic Antimicrobial Resistance Profiles From Whole Genome Sequences of Non-typhoidal Salmonella enterica, Saskia Neuert,

46. Whole-genome sequencing for prediction of Mycobacterium tuberculosis drug susceptibility and resistance: a retrospective cohort study, Timothy M Walker, Tim Walker's heuristic paper

47. VAMPr: VAriant Mapping and Prediction of antibiotic resistance via explainable features and machine learning, Jiwoong Kim, VAMPr paper

48. Use of whole genome sequencing for detection of antimicrobial resistance: Mycobacterium tuberculosis, a model organism., Vincent Escuyer, WGS for AMR - TB as a case study

#### 2 Data

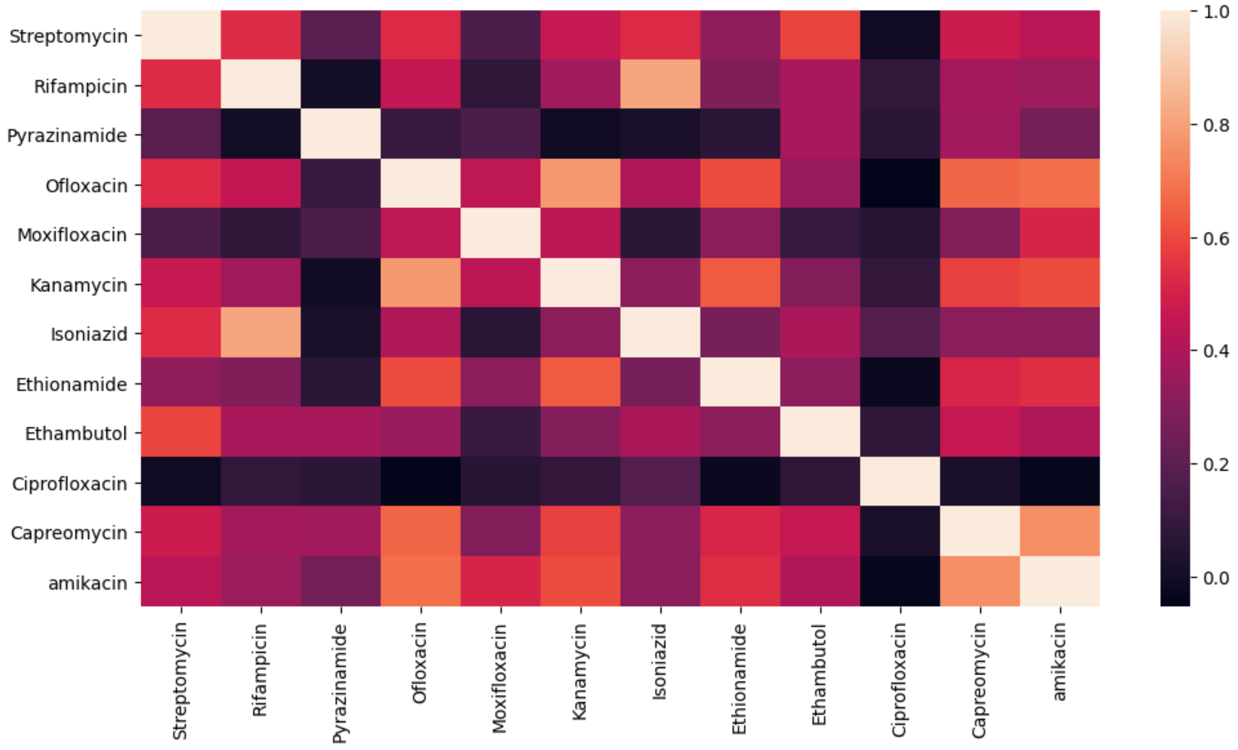


Figure 1: The pairwise correlation of the status vectors for each pair of drugs.

| | Streptomycin | Rifampicin | Pyrazinamide | Ofloxacin | Moxifloxacin | Kanamycin | Isoniazid | Ethionamide | Ethambutol | Ciprofloxacin | Capreomycin | amikacin |
|---|---|---|---|---|---|---|---|---|---|---|---|---|
| Streptomycin | 2104 | 1860 | 601 | 597 | 91 | 552 | 1989 | 342 | 1158 | 16 | 462 | 449 |
| Rifampicin | 1860 | 2968 | 667 | 750 | 117 | 654 | 2840 | 432 | 1342 | 35 | 522 | 542 |
| Pyrazinamide | 601 | 667 | 754 | 255 | 81 | 247 | 684 | 196 | 492 | 24 | 277 | 242 |
| Ofloxacin | 597 | 750 | 255 | 800 | 122 | 487 | 747 | 303 | 457 | 25 | 331 | 333 |
| Moxifloxacin | 91 | 117 | 81 | 122 | 129 | 85 | 113 | 67 | 51 | 18 | 66 | 74 |
| Kanamycin | 552 | 654 | 247 | 487 | 85 | 697 | 641 | 318 | 472 | 21 | 402 | 440 |
| Isoniazid | 1989 | 2840 | 684 | 747 | 113 | 641 | 3445 | 448 | 1378 | 31 | 512 | 530 |
| Ethionamide | 342 | 432 | 196 | 303 | 67 | 318 | 448 | 498 | 323 | 13 | 235 | 226 |
| Ethambutol | 1158 | 1342 | 492 | 457 | 51 | 472 | 1378 | 323 | 1407 | 22 | 402 | 399 |
| Ciprofloxacin | 16 | 35 | 24 | 25 | 18 | 21 | 31 | 13 | 22 | 37 | 12 | 8 |
| Capreomycin | 462 | 522 | 277 | 331 | 66 | 402 | 512 | 235 | 402 | 12 | 552 | 431 |
| amikacin | 449 | 542 | 242 | 333 | 74 | 440 | 530 | 226 | 399 | 8 | 431 | 573 |

Table 1: The pairwise similarity of the resistant status for each pair of drugs. Each cell represents the number of same samples between resistant isolates of that two drugs.

| | Streptomycin | Rifampicin | Pyrazinamide | Ofloxacin | Moxifloxacin | Kanamycin | Isoniazid | Ethionamide | Ethambutol | Ciprofloxacin | Capreomycin | amikacin |
|---|---|---|---|---|---|---|---|---|---|---|---|---|
| Streptomycin | 3021 | 2410 | 1525 | 1101 | 643 | 1104 | 2225 | 500 | 2651 | 85 | 657 | 726 |
| Rifampicin | 2410 | 4747 | 2651 | 893 | 506 | 844 | 4141 | 387 | 3608 | 212 | 545 | 614 |
| Pyrazinamide | 1525 | 2651 | 3104 | 708 | 401 | 509 | 2343 | 182 | 2820 | 214 | 673 | 594 |
| Ofloxacin | 1101 | 893 | 708 | 2111 | 731 | 1350 | 821 | 649 | 1321 | 48 | 1198 | 1237 |
| Moxifloxacin | 643 | 506 | 401 | 731 | 832 | 639 | 490 | 233 | 629 | 35 | 455 | 689 |
| Kanamycin | 1104 | 844 | 509 | 1350 | 639 | 1739 | 778 | 574 | 1206 | 187 | 915 | 970 |
| Isoniazid | 2225 | 4141 | 2343 | 821 | 490 | 778 | 4289 | 348 | 3208 | 59 | 512 | 554 |
| Ethionamide | 500 | 387 | 182 | 649 | 233 | 574 | 348 | 1018 | 605 | 26 | 454 | 481 |
| Ethambutol | 2651 | 3608 | 2820 | 1321 | 629 | 1206 | 3208 | 605 | 4689 | 354 | 872 | 893 |
| Ciprofloxacin | 85 | 212 | 214 | 48 | 35 | 187 | 59 | 26 | 354 | 406 | 89 | 60 |
| Capreomycin | 657 | 545 | 673 | 1198 | 455 | 915 | 512 | 454 | 872 | 89 | 1439 | 1113 |
| amikacin | 726 | 614 | 594 | 1237 | 689 | 970 | 554 | 481 | 893 | 60 | 1113 | 1460 |

Table 2: The pairwise similarity of the susceptible status for each pair of drugs. Each cell represents the number of same samples between susceptible isolates of that two drugs.

| Drug | Number of labelled isolates | Number of resistant isolates |
|---|---|---|
| Streptomycin | 1,136 | 87 (7.65%) |
| Rifampicin | 1,138 | 26 (2.28%) |
| Pyrazinamide | 134 | 13 (9.7%) |
| Isoniazid | 1,138 | 157 (13.79%) |
| Ethambutol | 1,138 | 22 (1.96%) |

Table 3: Summary of the number of isolates and the label distribution in BCCDC data.

### 3 Model parameters

| Layer \ Model | LRCN1 | LRCN2 | LRCN3 | LRCN4 | LRCN5 | LRCN6 | LRCN7 | LRCN8 | LRCN9 | LRCN10 |
|---|---|---|---|---|---|---|---|---|---|---|
| CNN1 (kernel, filter, pool) | (4, 5, 4) | (6, 8, 6) | (3, 8, 3) | (3, 8, 3) | (6, 8, 4) | (6, 8, 6) | (3, 8, 3) | (3, 8, 3) | (6, 8, 4) | (5, 7, 5) |
| CNN2 (kernel, filter, pool) | (5, 4, 4) | (3, 8, 3) | (6, 8, 6) | (4, 4, 4) | (3, 7, 3) | (6, 4, 4) | (6, 8, 6) | (6, 8, 4) | (6, 8, 6) | (4, 5, 4) |
| CNN3 (kernel, filter, pool) | (5, 7, 4) | - | (3, 7, 3) | - | - | (3, 4, 3) | (3, 4, 3) | (6, 4, 4) | (3, 8, 3) | (5, 7, 4) |
| CNN4 (kernel, filter, pool) | (4, 6, 4) | - | (3, 4, 3) | - | - | - | (3, 8, 3) | - | (3, 4, 3) | (6, 4, 4) |
| CNN5 (kernel, filter, pool) | - | - | (6, 8, 4) | - | - | - | (6, 4, 4) | - | (6, 8, 4) | - |
| LSTM1 | 478 | 64 | 64 | 439 | 478 | 457 | 64 | 413 | 518 | 518 |
| LSTM2 | 466 | 518 | 71 | 86 | 451 | 227 | 518 | 497 | 64 | 446 |
| LSTM3 | 350 | 518 | 518 | 518 | 460 | 350 | 454 | 518 | - | - |
| LSTM4 | 444 | 303 | 64 | 309 | 518 | - | - | 456 | - | - |
| LSTM5 | - | 518 | 64 | 518 | 518 | - | - | 518 | - | - |
| Dense1 | 306 | 518 | 518 | 159 | 356 | 518 | 64 | 515 | 518 | 277 |
| Dense2 | 293 | 64 | 64 | 518 | 416 | 518 | 95 | 362 | 467 | 76 |
| Dense3 | 325 | 64 | - | 518 | 286 | - | - | 518 | - | 348 |
| Dense4 | - | 64 | - | 64 | - | - | - | 77 | - | - |
| Dense5 | - | 64 | - | 317 | - | - | - | 415 | - | - |

Table 4: Parameters of LRCN models. Each row represents a layer and each column represents a model. The layers order in the table is the same of their order in the model.

| Layer \ Model | LSTM1 | LSTM2 | LSTM3 | LSTM4 | LSTM5 | LSTM6 | LSTM7 | LSTM8 | LSTM9 | LSTM10 |
|---|---|---|---|---|---|---|---|---|---|---|
| layer LSTM1 | 518 | 361 | 311 | 355 | 353 | 353 | 265 | 478 | 404 | 478 |
| layer LSTM2 | 64 | 142 | 401 | 453 | 237 | 237 | 116 | 466 | 482 | 237 |
| layer LSTM3 | - | 247 | 365 | 343 | 281 | 281 | 135 | 350 | - | 291 |
| layer LSTM4 | - | 291 | - | - | - | - | 313 | 444 | - | 444 |
| layer LSTM5 | - | - | - | - | - | - | - | - | - | 365 |
| Dense1 | 518 | 467 | 228 | 359 | 484 | 484 | 252 | 306 | 69 | 306 |
| Dense2 | 64 | 470 | 347 | 219 | 480 | 480 | 395 | 293 | 70 | 480 |
| Dense3 | 64 | 132 | 154 | 230 | 239 | 239 | 442 | 325 | 171 | - |
| Dense4 | 518 | 488 | 404 | 147 | - | - | 390 | - | 453 | - |
| Dense5 | 64 | - | 88 | 244 | - | - | - | - | 305 | - |

Table 5: Parameters of LSTM models. Each row represents a layer and each column represents a model. The layers order in the table is the same of their order in the model.

| Layer \ Model | WnD1 | WnD2 | WnD3 | WnD4 | WnD5 | WnD6 | WnD7 | WnD8 | WnD9 | WnD10 |
|---|---|---|---|---|---|---|---|---|---|---|
| Dense1 | 64 | 64 | 64 | 64 | 518 | 64 | 198 | 362 | 64 | 64 |
| Dense2 | 518 | 132 | 244 | 518 | 64 | 306 | 518 | 64 | 518 | 444 |
| Dense3 | 64 | 64 | 518 | - | - | 293 | 518 | 286 | - | 481 |
| Dense4 | 518 | 518 | 64 | - | - | - | 518 | 171 | - | 518 |
| Dense5 | 488 | 488 | 64 | - | - | - | 518 | 518 | - | 317 |
| kernel regularizer | 0.1 | 0.1 | 0.1 | 0.01 | 0.1 | 0.01 | 0.1 | 0.01 | 0.1 | 0.01 |

Table 6: Parameters of WnD models. Each row represents a layer and each column represents a model. The layers order in the table is the same of their order in the model.

| | RF1 | RF2 | RF3 | RF4 | RF5 | RF6 | RF7 | RF8 | RF9 | RF10 |
|---|---|---|---|---|---|---|---|---|---|---|
| n_estimators | 110 | 130 | 140 | 60 | 130 | 140 | 130 | 110 | 120 | 110 |
| min_samples_split | 4 | 4 | 4 | 4 | 4 | 3 | 4 | 4 | 4 | 3 |
| bootstrap | FALSE | FALSE | FALSE | FALSE | FALSE | FALSE | FALSE | FALSE | FALSE | FALSE |
| max_depth | 50 | 50 | None | 50 | 70 | 50 | 30 | 50 | None | 80 |

Table 7: Parameters of RF models. Each row represents a layer and each column represents a model.

| | LR1 | LR2 | LR3 | LR4 | LR5 | LR6 | LR7 | LR8 | LR9 | LR10 |
|---|---|---|---|---|---|---|---|---|---|---|
| C | 1 | 0.1 | 1 | 1 | 0.1 | 1 | 1 | 1 | 1 | 1 |
| max_iter | 657.7915 | 1000 | 657.7915 | 79.43 | 955.89 | 657.791473 | 657.791473 | 1000 | 1000 | 657.791473 |
| penalty | l2 | l1 | l1 | l1 | l2 | l1 | l1 | l2 | l1 | l1 |
| solver | newton-cg | liblinear | liblinear | liblinear | sag | liblinear | liblinear | newton-cg | liblinear | liblinear |

Table 8: Parameters of LR models. Each row represents a layer and each column represents a model.
The possible values for penalty were: l1 (solver = liblinear), l2 (solver = newton-cg, lbfgs, sag), elasticnet (solver = saga), none. The possible values for C were: (0.01, 0.1, 1, 10, 100)

| | SVM1 | SVM2 | SVM3 | SVM4 | SVM5 | SVM6 | SVM7 | SVM8 | SVM9 | SVM10 |
|---|---|---|---|---|---|---|---|---|---|---|
| C | 0.1 | 1 | 0.1 | 0.1 | 0.1 | 1 | 0.1 | 0.1 | 0.01 | 0.1 |
| kernel | linear | linear | linear | linear | linear | linear | linear | linear | linear | linear |
| gamma | - | - | - | - | - | - | - | - | - | - |
| degree | - | - | - | - | - | - | - | - | - | - |

Table 9: Parameters of SVM models. Each row represents a layer and each column represents a model.
The possible values for kernel were: linear, poly, rbf. The possible values for C were: (0.01, 0.1, 1, 10, 100)

| | GBT1 | GBT2 | GBT3 | GBT4 | GBT5 | GBT6 | GBT7 | GBT8 | GBT9 | GBT10 |
|---|---|---|---|---|---|---|---|---|---|---|
| max_depth | 130 | 140 | 90 | 60 | 70 | 90 | 50 | 60 | 50 | 70 |
| min_samples_split | 4 | 4 | 3 | 2 | 2 | 3 | 2 | 2 | 4 | 2 |
| n_estimators | 30 | 30 | 20 | 20 | 50 | 20 | 90 | 20 | 10 | 50 |
| random_state | 1 | 1 | - | 0 | - | - | 1 | 0 | - | - |

Table 10: Parameters of GBT models. Each row represents a layer and each column represents a model.

#### 3.1 KOVER

As mentioned before, KOVER produces discrete classifiers and therefore AUC-ROC and AUC-PR of this method can not be achieved at least in a traditional way. In this study, we calculated AUC-ROC using one point, and we were not able to calculate the sensitivity at 95% specificity and AUC-PR. To train KOVER models we used 10-fold cross-validation for hyper-parameter selection. We also set maximum number of rules in the model to 100 as opposed to 10 the default value in

KOVER (none of the trained models reached this limitation, since we tested other values as well to get the highest accuracy). Also the possible values for p were: 0.1, 1, 10, 100, 1000 and the chosen values by model were : 10, 1, 1, 10, 1, 1, 10, 1, 1, 10, 1

KOVER operates on the presence/absence of k-mers, therefore, we were not able to run it on non-binary gene burden features. Therefore, for training and evaluation of this method, we used SNP features that contain the presence/absence of each SNP in each isolate. As for other steps, we exactly applied them for KOVER as well.

### 4 Results

| | AUC-ROC | AUC-PR | sensitivity at 95% specificity | AUC-ROC on BCCDC |
|---|---|---|---|---|
| Streptomycin | 0.001 | 0.001 | 0.008 | 0.011 |
| Rifampicin | 0.001 | 0.001 | 0.001 | 0.009 |
| Pyrazinamide | 0.011 | 0.001 | 0.001 | 0.149 |
| Ofloxacin | 0.026 | 0.002 | 0.003 | - |
| Moxifloxacin | 0.058 | 0.189 | 0.001 | - |
| Kanamycin | 0.005 | 0.001 | 0.001 | - |
| Isoniazid | 0.001 | 0.001 (RF) | 0.001 | 0.002 |
| Ethionamide | 0.001 | 0.001 | 0.001 (WnD) | - |
| Ethambutol | 0.001 | 0.001 | 0.001 | 0.033 |
| Ciprofloxacin | 0.056 (GBT) | 0.006 (GBT) | 0.290 | - |
| Capreomycin | 0.001 | 0.001 | 0.005 | - |
| amikacin | 0.001 | 0.001 | 0.029 (WnD) | - |

Table 11: The p-values of experiments of Section 3.1 and 3.5. The names in parentheses shows the method which performs better than LRCN on that metric.

| | | Streptomycin | Rifampicin | Pyrazinamide | Ofloxacin | Moxifloxacin | Kanamycin | Isoniazid | Ethionamide | Ethambutol | Ciprofloxacin | Capreomycin | amikacin |
|---|---|---|---|---|---|---|---|---|---|---|---|---|---|
| LRCN | AUC-ROC | $0.918 \pm 0.003$ | $0.961 \pm 0.002$ | $0.911 \pm 0.006$ | $0.896 \pm 0.008$ | $0.880 \pm 0.022$ | $0.909 \pm 0.006$ | $0.931 \pm 0.002$ | $0.802 \pm 0.014$ | $0.911 \pm 0.005$ | $0.856 \pm 0.026$ | $0.865 \pm 0.010$ | $0.883 \pm 0.009$ |
| LRCN | AUC-PR | $0.861 \pm 0.004$ | $0.942 \pm 0.003$ | $0.749 \pm 0.015$ | $0.779 \pm 0.012$ | $0.633 \pm 0.045$ | $0.790 \pm 0.009$ | $0.892 \pm 0.004$ | $0.713 \pm 0.021$ | $0.746 \pm 0.010$ | $0.574 \pm 0.029$ | $0.736 \pm 0.018$ | $0.744 \pm 0.017$ |
| LRCN | R-95%-S | $0.508 \pm 0.050$ | $0.783 \pm 0.014$ | $0.544 \pm 0.021$ | $0.282 \pm 0.033$ | $0.484 \pm 0.033$ | $0.450 \pm 0.033$ | $0.629 \pm 0.063$ | $0.095 \pm 0.007$ | $0.543 \pm 0.025$ | $0.2 \pm 0.076$ | $0.284 \pm 0.029$ | $0.275 \pm 0.032$ |
| WnD | AUC-ROC | $0.832 \pm 0.007$ | $0.902 \pm 0.005$ | $0.858 \pm 0.007$ | $0.779 \pm 0.010$ | $0.766 \pm 0.018$ | $0.812 \pm 0.009$ | $0.864 \pm 0.003$ | $0.668 \pm 0.019$ | $0.852 \pm 0.006$ | $0.631 \pm 0.062$ | $0.743 \pm 0.011$ | $0.783 \pm 0.014$ |
| WnD | AUC-PR | $0.801 \pm 0.008$ | $0.853 \pm 0.009$ | $0.647 \pm 0.020$ | $0.537 \pm 0.023$ | $0.444 \pm 0.054$ | $0.643 \pm 0.021$ | $0.845 \pm 0.007$ | $0.520 \pm 0.034$ | $0.655 \pm 0.024$ | $0.186 \pm 0.049$ | $0.527 \pm 0.021$ | $0.588 \pm 0.020$ |
| WnD | R-95%-S | $0.430 \pm 0.043$ | $0.613 \pm 0.025$ | $0.445 \pm 0.032$ | $0.238 \pm 0.023$ | $0.316 \pm 0.046$ | $0.359 \pm 0.028$ | $0.548 \pm 0.019$ | $0.186 \pm 0.024$ | $0.473 \pm 0.024$ | $0.168 \pm 0.057$ | $0.245 \pm 0.025$ | $0.311 \pm 0.036$ |
| DeepAMR | AUC-ROC | $0.883 \pm 0.004$ | $0.927 \pm 0.003$ | $0.873 \pm 0.008$ | $0.662 \pm 0.011$ | $0.773 \pm 0.022$ | $0.790 \pm 0.013$ | $0.885 \pm 0.003$ | $0.639 \pm 0.012$ | $0.888 \pm 0.003$ | $0.602 \pm 0.070$ | $0.684 \pm 0.014$ | $0.703 \pm 0.015$ |
| DeepAMR | AUC-PR | $0.824 \pm 0.005$ | $0.898 \pm 0.004$ | $0.610 \pm 0.022$ | $0.342 \pm 0.018$ | $0.322 \pm 0.040$ | $0.579 \pm 0.036$ | $0.887 \pm 0.003$ | $0.396 \pm 0.016$ | $0.707 \pm 0.012$ | $0.200 \pm 0.082$ | $0.382 \pm 0.015$ | $0.404 \pm 0.015$ |
| DeepAMR | R-95%-S | $0.465 \pm 0.017$ | $0.659 \pm 0.016$ | $0.391 \pm 0.031$ | $0.017 \pm 0.005$ | $0.165 \pm 0.038$ | $0.274 \pm 0.031$ | $0.541 \pm 0.059$ | $0.052 \pm 0.014$ | $0.489 \pm 0.023$ | $0.035 \pm 0.035$ | $0.059 \pm 0.014$ | $0.068 \pm 0.014$ |
| LSTM | AUC-ROC | $0.830 \pm 0.010$ | $0.918 \pm 0.003$ | $0.872 \pm 0.013$ | $0.723 \pm 0.011$ | $0.827 \pm 0.028$ | $0.797 \pm 0.018$ | $0.825 \pm 0.004$ | $0.671 \pm 0.026$ | $0.828 \pm 0.007$ | $0.684 \pm 0.092$ | $0.745 \pm 0.031$ | $0.764 \pm 0.028$ |
| LSTM | AUC-PR | $0.797 \pm 0.010$ | $0.861 \pm 0.006$ | $0.669 \pm 0.022$ | $0.516 \pm 0.014$ | $0.525 \pm 0.051$ | $0.638 \pm 0.025$ | $0.824 \pm 0.008$ | $0.505 \pm 0.029$ | $0.620 \pm 0.022$ | $0.205 \pm 0.016$ | $0.567 \pm 0.026$ | $0.595 \pm 0.022$ |
| LSTM | R-95%-S | $0.301 \pm 0.027$ | $0.353 \pm 0.007$ | $0.308 \pm 0.016$ | $0.171 \pm 0.017$ | $0.314 \pm 0.037$ | $0.251 \pm 0.021$ | $0.320 \pm 0.006$ | $0.114 \pm 0.017$ | $0.304 \pm 0.014$ | $0.083 \pm 0.083$ | $0.197 \pm 0.019$ | $0.194 \pm 0.022$ |
| GBT | AUC-ROC | $0.898 \pm 0.004$ | $0.953 \pm 0.003$ | $0.895 \pm 0.006$ | $0.889 \pm 0.007$ | $0.860 \pm 0.023$ | $0.899 \pm 0.007$ | $0.909 \pm 0.003$ | $0.763 \pm 0.013$ | $0.892 \pm 0.006$ | $0.886 \pm 0.040$ | $0.837 \pm 0.012$ | $0.852 \pm 0.009$ |
| GBT | AUC-PR | $0.800 \pm 0.005$ | $0.872 \pm 0.002$ | $0.721 \pm 0.015$ | $0.753 \pm 0.019$ | $0.601 \pm 0.040$ | $0.771 \pm 0.013$ | $0.881 \pm 0.004$ | $0.650 \pm 0.026$ | $0.722 \pm 0.015$ | $0.654 \pm 0.076$ | $0.701 \pm 0.018$ | $0.717 \pm 0.020$ |
| GBT | R-95%-S | $0.460 \pm 0.011$ | $0.535 \pm 0.006$ | $0.469 \pm 0.052$ | $0.192 \pm 0.019$ | $0.401 \pm 0.053$ | $0.340 \pm 0.023$ | $0.513 \pm 0.006$ | $0.201 \pm 0.026$ | $0.360 \pm 0.021$ | $0.083 \pm 0.112$ | $0.147 \pm 0.055$ | $0.171 \pm 0.036$ |
| KOVER | AUC-ROC | $0.850 \pm 0.000$ | $0.928 \pm 0.000$ | $0.748 \pm 0.000$ | $0.890 \pm 0.000$ | $0.796 \pm 0.000$ | $0.829 \pm 0.000$ | $0.908 \pm 0.000$ | $0.777 \pm 0.000$ | $0.708 \pm 0.000$ | $0.780 \pm 0.000$ | $0.817 \pm 0.000$ | $0.854 \pm 0.000$ |
| KOVER | AUC-PR | $0.000 \pm 0.000$ | $0.000 \pm 0.000$ | $0.000 \pm 0.000$ | $0.000 \pm 0.000$ | $0.000 \pm 0.000$ | $0.000 \pm 0.000$ | $0.000 \pm 0.000$ | $0.000 \pm 0.000$ | $0.000 \pm 0.000$ | $0.000 \pm 0.000$ | $0.000 \pm 0.000$ | $0.000 \pm 0.000$ |
| KOVER | R-95%-S | $0.000 \pm 0.000$ | $0.000 \pm 0.000$ | $0.000 \pm 0.000$ | $0.000 \pm 0.000$ | $0.000 \pm 0.000$ | $0.000 \pm 0.000$ | $0.000 \pm 0.000$ | $0.000 \pm 0.000$ | $0.000 \pm 0.000$ | $0.000 \pm 0.000$ | $0.000 \pm 0.000$ | $0.000 \pm 0.000$ |
| RF | AUC-ROC | $0.884 \pm 0.004$ | $0.947 \pm 0.002$ | $0.896 \pm 0.008$ | $0.797 \pm 0.008$ | $0.854 \pm 0.013$ | $0.851 \pm 0.010$ | $0.895 \pm 0.004$ | $0.675 \pm 0.008$ | $0.885 \pm 0.003$ | $0.660 \pm 0.083$ | $0.780 \pm 0.009$ | $0.796 \pm 0.011$ |
| RF | AUC-PR | $0.831 \pm 0.006$ | $0.925 \pm 0.003$ | $0.711 \pm 0.015$ | $0.586 \pm 0.029$ | $0.562 \pm 0.036$ | $0.714 \pm 0.023$ | $0.895 \pm 0.003$ | $0.451 \pm 0.016$ | $0.670 \pm 0.019$ | $0.359 \pm 0.095$ | $0.587 \pm 0.023$ | $0.608 \pm 0.024$ |
| RF | R-95%-S | $0.069 \pm 0.069$ | $0.489 \pm 0.133$ | $0.258 \pm 0.106$ | $0.117 \pm 0.078$ | $0.0 \pm 0.0$ | $0.129 \pm 0.086$ | $0.313 \pm 0.128$ | $0.095 \pm 0.064$ | $0.236 \pm 0.096$ | $0.0 \pm 0.0$ | $0.227 \pm 0.094$ | $0.188 \pm 0.097$ |
| LR | AUC-ROC | $0.882 \pm 0.004$ | $0.943 \pm 0.003$ | $0.904 \pm 0.004$ | $0.887 \pm 0.004$ | $0.839 \pm 0.012$ | $0.876 \pm 0.008$ | $0.898 \pm 0.005$ | $0.754 \pm 0.016$ | $0.877 \pm 0.004$ | $0.762 \pm 0.051$ | $0.823 \pm 0.007$ | $0.840 \pm 0.012$ |
| LR | AUC-PR | $0.825 \pm 0.006$ | $0.879 \pm 0.003$ | $0.720 \pm 0.009$ | $0.752 \pm 0.011$ | $0.590 \pm 0.040$ | $0.759 \pm 0.010$ | $0.857 \pm 0.005$ | $0.654 \pm 0.022$ | $0.709 \pm 0.009$ | $0.309 \pm 0.112$ | $0.669 \pm 0.013$ | $0.687 \pm 0.017$ |
| LR | R-95%-S | $0.0 \pm 0.0$ | $0.0 \pm 0.0$ | $0.257 \pm 0.105$ | $0.0 \pm 0.0$ | $0.086 \pm 0.058$ | $0.055 \pm 0.055$ | $0.0 \pm 0.0$ | $0.0 \pm 0.0$ | $0.064 \pm 0.064$ | $0.1 \pm 0.1$ | $0.0 \pm 0.0$ | $0.0 \pm 0.0$ |
| SVM | AUC-ROC | $0.865 \pm 0.006$ | $0.932 \pm 0.002$ | $0.888 \pm 0.006$ | $0.881 \pm 0.006$ | $0.830 \pm 0.016$ | $0.864 \pm 0.006$ | $0.886 \pm 0.003$ | $0.750 \pm 0.013$ | $0.869 \pm 0.005$ | $0.818 \pm 0.043$ | $0.817 \pm 0.008$ | $0.823 \pm 0.009$ |
| SVM | AUC-PR | $0.815 \pm 0.007$ | $0.867 \pm 0.003$ | $0.705 \pm 0.013$ | $0.759 \pm 0.012$ | $0.547 \pm 0.033$ | $0.738 \pm 0.010$ | $0.841 \pm 0.004$ | $0.644 \pm 0.023$ | $0.712 \pm 0.010$ | $0.369 \pm 0.071$ | $0.669 \pm 0.015$ | $0.672 \pm 0.010$ |
| SVM | R-95%-S | $0.0 \pm 0.0$ | $0.0 \pm 0.0$ | $0.137 \pm 0.091$ | $0.0 \pm 0.0$ | $0.078 + -0.053$ | $0.0 \pm 0.0$ | $0.0 \pm 0.0$ | $0.0 \pm 0.0$ | $0.0 \pm 0.0$ | $0.05 \pm 0.05$ | $0.0 \pm 0.0$ | $0.0 \pm 0.0$ |

Table 12: The performance and confidence intervals of all models in Figure 3.
